# Tuning the brain at 40 Hz: Synergistic effects of combined tACS and auditory steady-state response

**DOI:** 10.64898/2026.04.24.720765

**Authors:** Fabio Masina, Rachele Pezzetta, Nicole Genero, Alessandro Tonin, Giorgio Arcara, Daniela Mapelli

## Abstract

Gamma oscillations are altered in several conditions such as schizophrenia, Alzheimer’s Disease, and Mild Cognitive Impairment. Both 40-Hz transcranial alternating current stimulation (tACS) and 40-Hz auditory steady-state stimulation can entrain gamma oscillations and improve cognitive outcomes, but their combined effects remain unclear. This study tested whether preconditioning gamma activity with tACS enhances the auditory steady-state response (ASSR) to 40-Hz auditory stimulation, reflecting a potential synergistic interaction.

In a within-subject, sham-controlled design, EEG was recorded before and after 40-Hz tACS delivered over bilateral sensorimotor areas in 34 healthy participants. 40-Hz auditory stimulation was administered before and after tACS to evaluate potential changes in ASSR. Source-level gamma power was analyzed in temporal and sensorimotor regions using linear mixed-effects models.

Compared to sham, real tACS increased ASSR-related gamma activity in the superior temporal gyrus, with no effects elsewhere or between hemispheres. These findings support the use of multimodal gamma entrainment to enhance neural oscillations.

To conclude, combining 40-Hz tACS with auditory stimulation enhances gamma activity more than either intervention alone, supporting potential clinical applications for neuropsychiatric conditions with disrupted gamma oscillations.

## 1. Introduction

Brain gamma oscillations range approximately from 30 to 100 Hz and are thought to arise from the interplay between inhibitory gamma-aminobutyric acid (GABA)ergic interneurons and excitatory inputs from pyramidal cells ^1^. Gamma oscillations exert a crucial role in cognitive processing, since they are elicited by several higher-order cognitive functions, including memory and attention ^2^. Notably, electrophysiological studies show that gamma band synchronization increases during attentional selection ^3^ and is sustained during the maintenance of short-term and working memory representations ^4^. Moreover, enhanced gamma activity during stimulus encoding has been shown to predict successful long-term memory formation ^5,6^.

Neurodegenerative diseases can involve a prominent disruption in gamma oscillations. As a matter of fact, aberrant gamma oscillatory activity has been shown in clinical conditions such as Alzheimer’s Disease (AD) or Mild Cognitive Impairment (MCI) ^7–9^. Within the AD framework, the alteration of gamma usually falls within the 38-42 Hz band, a frequency range closely linked to hippocampal mechanisms underlying memory processing ^10^. Clinically, this disruption is proportional to the advancement of AD ^7,11^ and is functionally linked to worse memory performance, both in AD and amnestic MCI patients ^12,13^.

Given the central role of gamma oscillations in AD pathology, research has been prompted into whether restoring this rhythm could have therapeutic effects for these patients ^14^. Indeed, a landmark study from Iaccarino et al., ^15^ demonstrated that entraining gamma oscillations reduced AD-related pathology in mice, an effect replicated across multiple paradigms ^16,17^. Furthermore, restoration of gamma oscillation amplitude was sufficient to recover spatial memory in mice ^18^.

Driven by these findings, researchers are investigating methods to entrain human gamma oscillations, with transcranial alternating current stimulation (tACS) emerging as a leading technique. tACS delivers oscillating currents ^19^ via two or more electrodes placed on the scalp, generating rhythmic electric fields in the brain. Rather than directly altering neuronal excitability, tACS interacts with ongoing neural dynamics, thereby promoting the entrainment and synchronization of oscillatory brain networks in a frequency- and phase-specific manner ^20,21^. Beyond transient online effects, growing evidence indicates that tACS can also induce post-stimulation changes in synaptic plasticity or network connectivity ^22–24^.

A growing body of work has begun to target gamma oscillations, thereby setting tACS in the 30-100 Hz range ^25^. Gamma-tACS has been proven to ameliorate episodic memory processing in both AD and MCI patients ^26–29^. Findings from Benussi et al., ^26^ were later confirmed in a follow-up ^30^, with EEG showing specific gamma entrainment following active 40-Hz tACS. Taken together, these studies suggest that gamma-tACS is able to efficiently and selectively modulate memory-relevant gamma-band activity, potentially paving the way for novel therapeutic interventions ^31^.

Beyond direct entrainment through gamma-tACS, sensory stimulation with oscillatory patterns provides another method to modulate gamma oscillations through exogenous rhythmic inputs at 40 Hz (for a review, see Herrmann et al. ^32^). This type of stimulation can evoke a 40 Hz steady-state response (SSR), indicating early sensory activity that occurs prior to higher-order perceptual and attentional processing ^33^. Entraining gamma oscillations through repetitive auditory stimulation has been deemed successful in healthy human participants ^34^. Notably, the auditory steady-state response (ASSR) at 40 Hz reflects the phase- and frequency-locking of EEG oscillations to periodic auditory stimuli such as amplitude-modulated tones or click trains with a fixed inter-click interval ^35^. This evoked response mainly originates from the auditory cortex and its interconnected neural pathways ^36^, although several additional brain regions have been identified as neural generators beyond the auditory cortex ^37^.

Disruptions in the ASSR have been reported across several clinical conditions. This phenomenon translates into measurable alterations of response amplitude and phase stability, suggesting impaired neural synchronization ^35^. Specifically, the ASSR has been previously identified as a potential biomarker for schizophrenia ^35,38^, in which several neural oscillations associated with specific neural circuits are impaired ^39^, and it is compromised in people with AD ^5,33^ and MCI ^40^. In addition, the brain’s capacity to generate synchronized gamma activity in response to 40-Hz periodic auditory stimuli appears to serve as a marker of rehabilitation success in stroke patients ^41^.

Beyond its potential role as a biomarker of schizophrenia, AD, MCI, and functional recovery, the ASSR may also exert beneficial, intervention-like effects, similar to those reported for gamma-tACS. Entrainment of gamma activity through periodic auditory stimulation has been associated with therapeutic benefits in conditions characterized by disrupted gamma oscillations, including schizophrenia ^39^, AD, and MCI ^15^.

Taken together, current evidence suggests that both gamma-tACS and 40-Hz auditory stimulation can induce gamma-frequency entrainment. Nevertheless, no evidence currently suggests that combining these approaches produces an additional (i.e., synergistic) increase in gamma activity. In the present study, we combined gamma-tACS with 40-Hz auditory stimulation to test whether inducing a gamma oscillatory state with tACS would potentiate the ASSR. Our hypothesis was that gamma-tACS would first tune cortical gamma oscillations, such that subsequent 40-Hz auditory stimulation, in line with a state-dependent framework ^42^, would interact synergistically with this pre-modulated activity and allow the ASSR to emerge with greater gamma power than auditory stimulation alone (i.e., the control condition with sham).

Studying healthy participants allows us to test the feasibility and state-dependent effects of combined exogenous (i.e., auditory and electrical) gamma-frequency stimulations, providing a foundation for future research in patient populations. In this study, we therefore examined the effects of combined gamma-tACS and 40-Hz auditory stimulation in a cohort of healthy adults to test the feasibility and state-dependent enhancement of gamma-band synchronization. In particular, we implemented a within-subject, counterbalanced, sham-controlled design that comprised three conditions: a pre-stimulation ASSR measurement via EEG, followed by gamma-tACS (real and sham), and a final post-stimulation ASSR measurement.

## 2. Materials and Methods

### 2.1. Participants

The study was conducted in agreement with the 1964 Helsinki Declaration and received approval from the local ethics committee (Institutional Ethics Committee Approval No. 2021.18). Participants were informed about the aims, procedures, potential risks and benefits of the research, and subsequently provided written informed consent. Data collection took place between March 9, 2022 and September 21, 2022.

A power analysis was carried out with G*Power 3 program ^43^ to determine the appropriate sample size. For the present study, assuming a medium effect size (*f* = 0.25), an alpha of 0.05, and a power of 0.9, the minimum required number of participants was estimated to be 30.

Thirty-four healthy volunteers participated in the study (22 females; mean age = 28.4, *SD* ± 4.2; mean education = 19.4 years, *SD* ± 1.7), and none of them reported any history of neurological or psychiatric disorders. Participants were primarily recruited among university students and hospital staff. No monetary or other incentives were provided, apart from reimbursement of travel expenses. All volunteers were screened for tACS exclusion criteria^44^. Of the 34 participants, 32 were right-handed according to Oldfield’s Edinburgh Handedness Inventory ^45^. This study complies with the Preferred Evaluation of Cognitive and Neuropsychological Studies (PECANS) guidelines ^46^. The completed PECANS checklist is provided in the Supplementary Materials.

### 2.2. Procedure and ASSR paradigm

Data collection took place in 2024 as part of a broader investigation. The original study ^47^ focused on the exploration of tACS modulation mechanisms over the aperiodic EEG component, examining hypotheses that are unrelated to the current research questions. The present study represents an independent secondary analysis of that dataset and tests different hypotheses related to potential synergistic effects of the application of gamma-tACS and 40-Hz auditory stimulation.

A paired, counterbalanced, sham-controlled design was adopted. It included two stimulation sessions, held on different days, during which bilateral tACS was applied. Each participant underwent both real and sham tACS in a counterbalanced sequence, with at least five days between sessions to reduce possible carry-over effects. Each session comprised three phases (*Figure 1*). In the initial phase (pre-stimulation), EEG data were collected during a 40-Hz auditory stimulation (i.e., the ASSR paradigm) triggering the ASSR. The second phase (stimulation) consisted of a 20-minute stimulating session during which either real or sham tACS was applied. Participants remained quietly seated throughout the stimulation phase. The final phase (post-stimulation) mirrored the pre-stimulation phase, repeating the ASSR paradigm.

**Fig 1.**
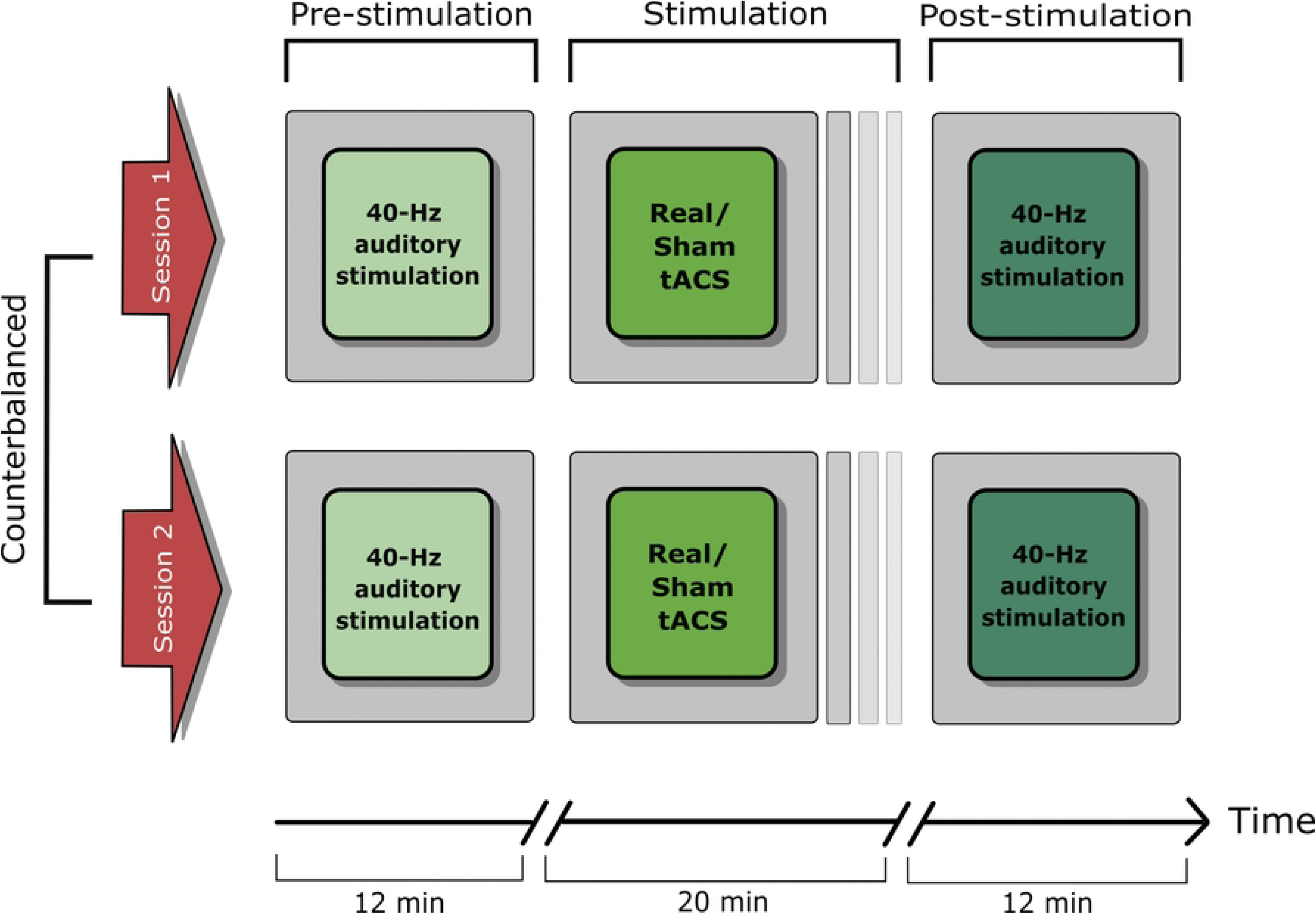
Representation of the experimental procedure. In each condition (real vs sham) participants underwent a both pre-stimulation and post-stimulation phases, during which 40-Hz auditory stimulation was applied. Real or sham tACS was applied in between those two phases, for about 20 min. EEG was recorded in both the pre-and post-stimulation phases.

During both the pre-stimulation and post-stimulation phases, participants underwent EEG recording with their eyes open and positioned 60 cm away from a computer screen. Throughout the EEG recording, participants were instructed to maintain their gaze on a fixation point (i.e., a cross in the middle of the screen).

The ASSR paradigm was created using the PsychoPy3 Experiment Builder (version 3.06; Peirce ^48,49^) and delivered both before and after the tACS session. The stimuli consisted of 40-Hz click trains lasting 2 seconds and played binaurally at 3 dB ^50^. The ASSR paradigm consisted of 180 trials, with every trial containing an auditory stimulus followed by a 2-second silent interval (ISI = 2 seconds). A white fixation cross remained visible on the screen for the entire duration of the task, during both stimulation and silent intervals. Participants sat comfortably facing the monitor throughout all the duration of the experimental phase.

### 2.3. Transcranial alternating current stimulation

During the stimulation phase, tACS was applied in accordance with the updated guidelines provided by Antal et al., ^44^. Two circular sponge electrodes soaked in saline solution (surface area: 8 cm²; current density: 0.25 mA/cm²) were positioned over the sensorimotor (SM) areas.

Prior to data collection, we conducted a finite element method simulation using SimNIBS ^51^, which is designed to measure the spatial distribution and strength of electric fields produced by both transcranial magnetic and transcranial electrical stimulation. This modeling indicated that the electric field generated by our tACS setup was primarily concentrated over the SM regions (*Figure 2*).

**Fig 2.**
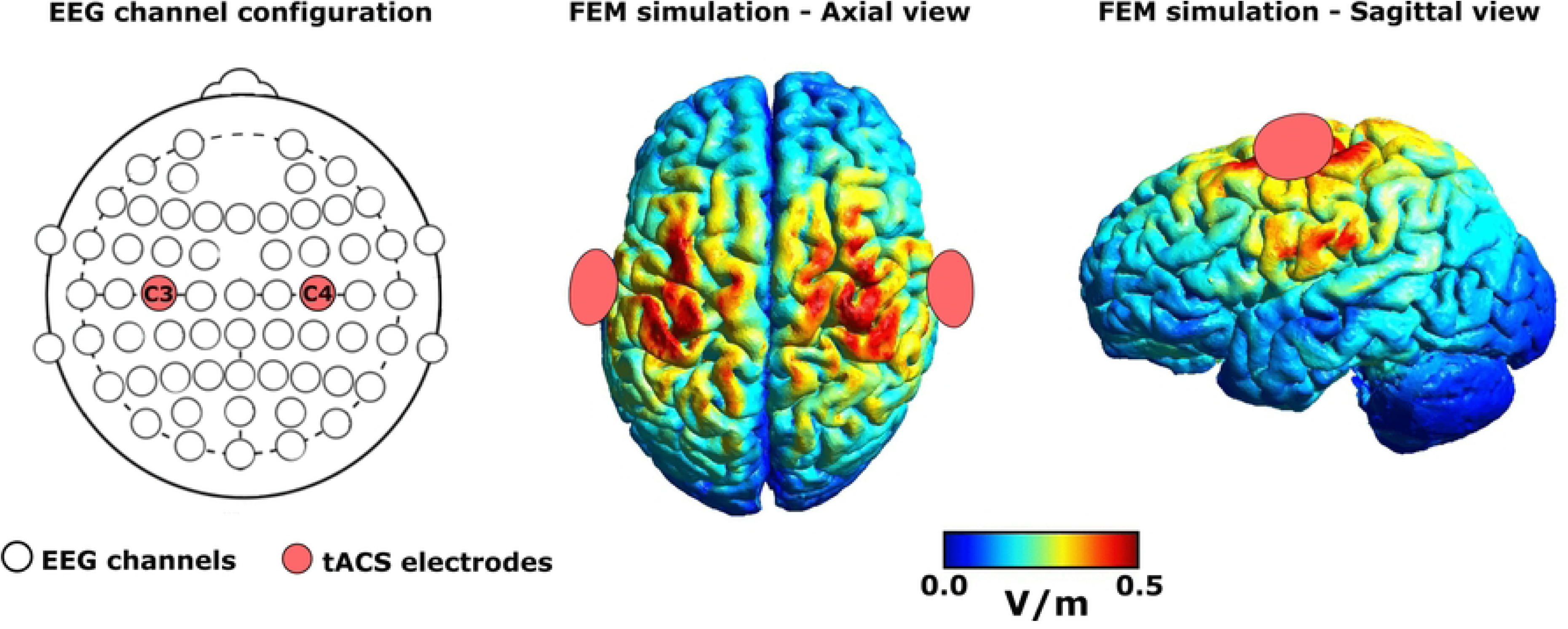
EEG and tACS configurations. The EEG channel configuration is displayed on the left. The two tACS electrodes were positioned over C3 and C4. The middle and right images illustrate the tACS-induced electric field distribution, estimated using SimNIBS through the finite element method ^51^

We selected the bilateral SM cortices (corresponding to C3 and C4 of the international 10–20 EEG system) as stimulation targets, despite the ASSR being maximal in auditory regions ^52^, based on evidence indicating strong anatomical and functional interconnections between SM areas and auditory cortices ^53^. These SM regions are part of distributed cortical networks that can influence auditory processing and oscillatory synchronization ^54,55^, which aligns with our expectations in the present study. Notably, previous neuromodulation studies have demonstrated that stimulation of SM cortices using tDCS can modulate gamma synchronization in temporal auditory regions, despite the absence of direct stimulation of the auditory cortex ^53^.

In addition, recent evidence indicates that tACS efficacy depends on electrode montage and the resulting field strength ^56^. Because the auditory cortex is relatively deep, direct stimulation may be less effective there, whereas more superficial and accessible regions like the SM cortices allow stronger and more consistent modulation ^53^. This supports targeting SM cortices as effective upstream sites for modulating auditory gamma activity with tACS.

Stimulation was delivered using a battery-powered device (BrainStim, EMS Medical, Italy). The tACS electrodes were mounted beneath the EEG cap at positions C4 and C3 of the international 10–20 system, corresponding to the right and left SM areas, respectively ^57,58^ . The tACS electrodes remained in place throughout the experimental session.

The stimulation protocol adopted a frequency of 40 Hz and a peak-to-peak intensity of 2 mA. Both real and sham conditions featured 30-second ramp-up and ramp-down periods. Active stimulation lasted for 20 minutes, however, in the sham condition no current was applied apart from the brief ramp periods at the beginning and end of the expected stimulation interval. After each session, participants were asked to answer a questionnaire assessing any sensations associated with the transcranial electrical stimulation ^59^. Importantly, participants were unable to reliably identify whether they received real or sham tACS [Session 1: Wald χ²(1) = 0.35, *p* = 0.551; Session 2: Wald χ²(1) = 0.28, *p* = 0.594].

### 2.4. EEG acquisition and data preprocessing

EEG was recorded in each session with a sampling rate of 1000 Hz using an actiCHamp amplifier equipped with 64 active electrodes (Brain Products GmbH, Germany). Recordings were referenced online to AFz, with FPz serving as the ground. Electrode impedance was maintained below 5 kΩ for the entire duration of the session.

Offline preprocessing was conducted with EEGLAB v2024.1 running on Matlab R2021b (The MathWorks Natick, MA, USA). The first step consisted of removing channels C3, C4, Iz and one external channel, yielding a total of 60 channels for analysis. Channels C3 and C4 were excluded because they were occupied by the tACS electrodes during recording. The remaining data were then down-sampled to 500 Hz. Subsequently, a band-pass filter from 0.1 to 70 Hz was applied and 50-Hz line noise was eliminated using the CleanLine plugin in EEGLAB (https://www.nitrc.org/projects/cleanline/).

After filtering, the continuous data were visually inspected to discard any channels exhibiting excessive noise. Residual artifacts, including ocular activity and muscle-related components, were identified and then removed by applying independent component analysis ^60^. The independent components were inspected using scalp topography, spectral features, temporal characteristics, and amplitude ^61^. The cleaned data were segmented into 5-second epochs and re-referenced to the average signal of each epoch. Epochs with amplitude exceeding ±100 µV were automatically removed. Further details on the EEG preprocessing are provided in Table 1.

**Table 1.** Details on the EEG preprocessing during the ASSR paradigm. Rejected channels and rejected Independent Components (ICs) are reported as means and standard deviations (*SDs*).

|  | Pre-stimulation phase |  | Post-stimulation phase |  |
| --- | --- | --- | --- | --- |
|  | Real | Sham | Real | Sham |
| Rejected channels | 2 (2.2) | 3 (2.8) | 1 (1.5) | 1 (2) |
| Rejected ICs | 9 (9) | 10 (11.4) | 8 (7.3) | 8 (8.3) |
| Rejected epochs | 13.9% | 18.9% | 8.8% | 19.5% |
| Epoch length | 5 s | 5 s | 5 s | 5 s |

### 2.5. Source estimation and ASSR data analysis

To test the hypothesis that combining gamma-tACS and 40-Hz auditory stimulation would produce synergistic increase in gamma activity, four clusters of regions of interest (ROIs) were selected a priori based on their known involvement in the ASSR generation and sensory processing. Specifically, the four ROIs were defined to capture both primary auditory ASSR sources and sensorimotor regions directly targeted by stimulation. The superior temporal gyrus (STG) and superior temporal sulcus (STS) were selected due to their central role in auditory processing ^62^ and their robust involvement in the ASSR generation during 40-Hz auditory stimulation ^52^. In addition, the precentral and postcentral gyri were included to represent primary motor and somatosensory cortices, respectively, allowing us to assess potential propagation of gamma modulation within interconnected SM and auditory networks.

Source estimation was performed on the preprocessed EEG data using sLORETA (constrained solution, 2018 implementation), without explicit noise modeling. Cleaned EEG data were segmented into 5-s epochs (−2500 to +2500 ms relative to stimulus onset) and projected to source space. Time–frequency decomposition was conducted at the source level using complex Morlet wavelets. Wavelet parameters were optimized to target gamma-band activity, with a central frequency of 40 Hz (MorletFc = 40) and a full width at half maximum (FWHM) of 0.5. Power estimates were computed as magnitude values and normalized using z-score normalization relative to a pre-stimulus baseline (−250 to −50 ms). Regional time series were extracted using the Desikan–Killiany cortical atlas ^63^, selecting bilateral auditory and sensorimotor scouts. The following anatomical regions were included: *banks of the superior temporal sulcus* (bankssts L/R), *superior temporal gyrus* (superiortemporal L/R), *postcentral gyrus* (postcentral L/R), and *precentral gyrus* (precentral L/R). Within each ROI, source activity was summarized using a principal component analysis (PCA) scout function, retaining the dominant component as representative of regional activation.

For ASSR-specific analyses, gamma-band power was obtained by averaging source-level activity within the 39–41 Hz frequency range ^53^. Based on the temporal dynamics of the ASSR, which reached its maximum at 250 ms and continues until the end of the stimulus ^64^, mean gamma power was extracted from a post-stimulus time window of 300–400 ms, corresponding to the period in which the steady-state response is fully established and stable^65^. This analysis pipeline enabled a targeted evaluation of gamma-band entrainment at the cortical source level and allowed testing for synergistic effects of concurrent gamma-tACS and 40-Hz auditory stimulation across auditory and SM networks.

Outlier values were removed prior to statistical analysis. Specifically, values exceeding ±2 standard deviations from both pre- and post-stimulation gamma power were excluded. This approach minimized the influence of extreme values while preserving within-ROI variability relevant to condition-dependent effects.

### 2.6. Statistical analysis

All data were analyzed using RStudio software (version 2025.05.1) and package lme4 ^66^ and lmerTest ^67^. All relevant data and R scripts are available at the OSF link (https://osf.io/gmq34/overview?view_only=a190d4432e7842fca4050b20ee8b499d). To test our hypothesis, linear mixed-effects models were conducted. In the statistical models, post-stimulation EEG measurements were adjusted for baseline levels as recommended ^68^ and implemented in previous studies ^57,69,70^. In all the models, the dependent variable was gamma power in the post-stimulation phase. The fixed effects terms were assessed by means of *F*-test using Satterthwaite approximation ^71^. To further characterize significant effects, post-hoc pairwise comparisons were performed using estimated marginal means, with *p*-values adjusted for multiple comparisons using the Tukey method.

In the main model, ROI (STS, STG, postcentral and precentral gyri), Stimulation (real vs. sham tACS), and their interaction (ROI x Stimulation) were entered as fixed effects. In addition, gamma power in the pre-stimulation phase was included as a covariate to adjust for baseline differences. The random-effect structure also included a random intercept for participants.

In addition, in the case this first model showed a significant ROI × Stimulation interaction, follow-up models were conducted on each specific ROI to further explore hemispheric differences. These models included Laterality (left vs. right), Stimulation (real vs. sham tACS), and their interaction (Laterality x Stimulation).

## 3. Results

The 40-Hz auditory stimulation elicited the typical ASSR, as reflected by a post-stimulus increase in gamma-band activity in the auditory cortex (*Figure 3*).

**Fig 3.**
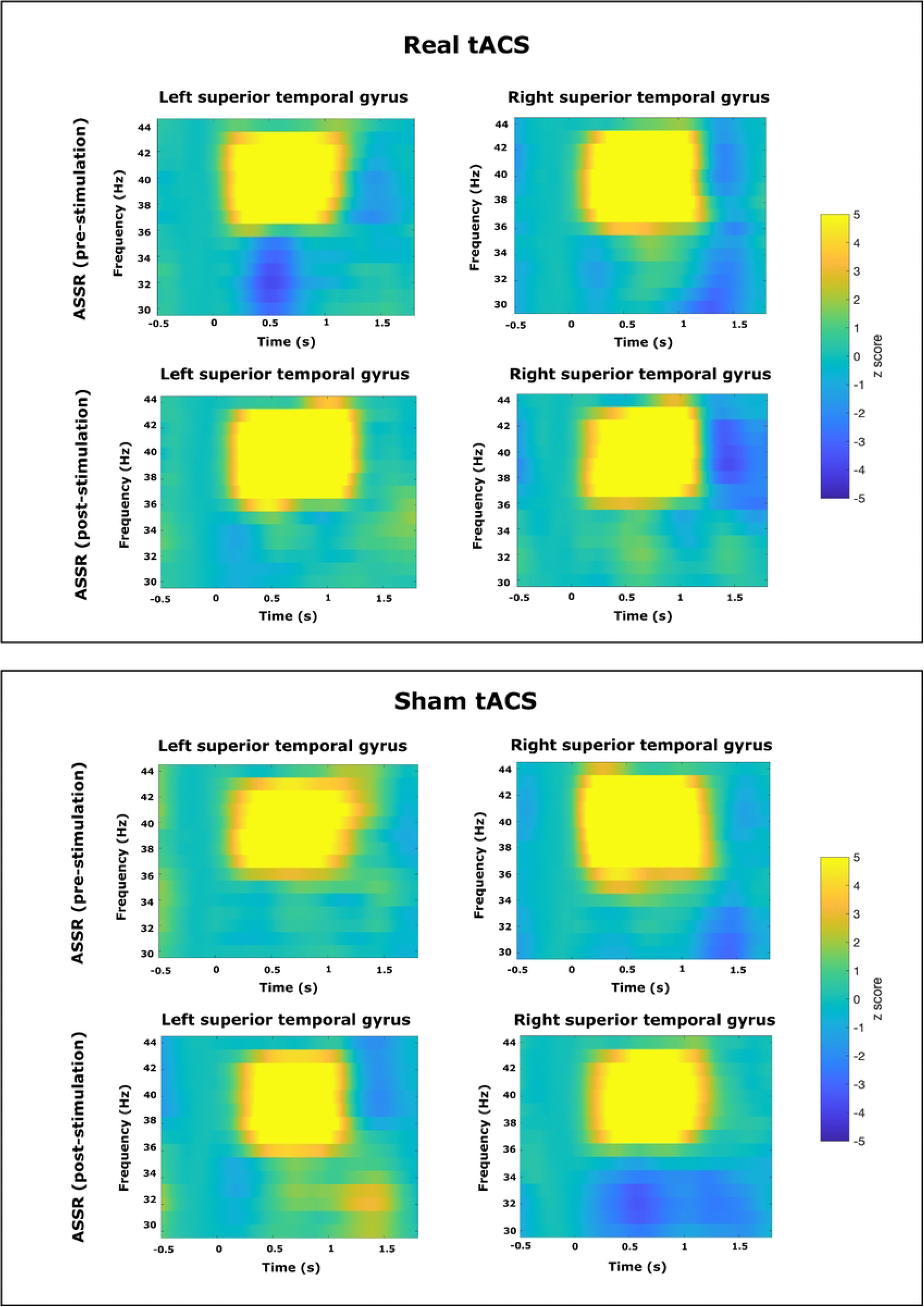
ASSR representation in the STG. Time-frequency representations of the ASSR in the left and right STG during real (top panel) and sham (bottom panel) tACS. For each condition, the ASSR is shown separately for pre-stimulation (top row) and post-stimulation (bottom row). Color values represent z-scored power relative to baseline across frequencies (30-45 Hz) and time

The main model revealed a significant main effect of ROI (*F*(3,456.55) = 11.38, *p* < .001), indicating differences in gamma power across cortical regions. Post-hoc pairwise comparisons for the main effect of ROI (Tukey-corrected) revealed that gamma power was significantly higher in the STS compared to both the Postcentral (estimate = 5.67, SE = 1.53, *p* = .0013) and Precentral regions (estimate = 7.79, SE = 1.54, *p* < .001). Similarly, the STG showed significantly greater gamma power than both the Postcentral (estimate = 4.42, SE = 1.53, *p* = .0204) and Precentral regions (estimate = 6.53, SE = 1.53, *p* < .001). In contrast, no significant differences were observed between STS and STG (*p* = .85), nor between Postcentral and Precentral regions (*p* = .50). No significant main effect of Stimulation was observed (*F*(1,460.13) = 0.74, *p* = .39).

Crucially, the interaction between ROI and Stimulation was significant (*F*(3,456.33) = 3.19, *p* = .024), suggesting that the effects of stimulation varied across regions. Post-hoc pairwise comparisons (Tukey-corrected) showed that real tACS was associated with significantly higher gamma power compared to sham tACS over the STG (estimate = 6.71, SE = 2.20, *p* =.048). In contrast, no significant differences between real and sham stimulation were observed in the STS, postcentral gyrus or precentral gyrus (all *p* >.05).

Figure 4 illustrates individual post-stimulation gamma power values for real and sham tACS across the analyzed ROIs, highlighting the higher gamma power during real tACS in the STG. In addition, the distribution of gamma power values across conditions and regions is illustrated in Figure 5.

**Fig 4.**
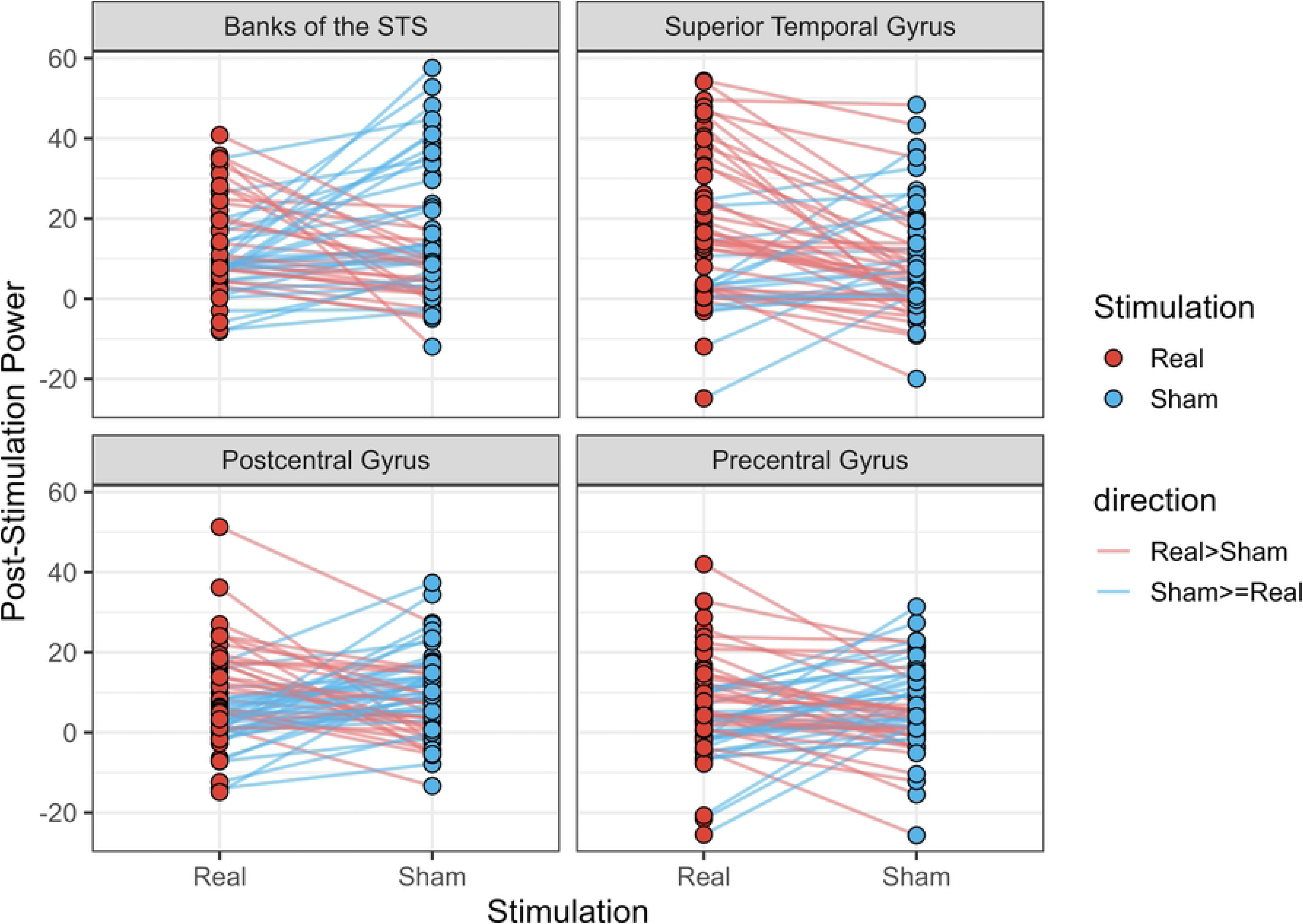
Individual trajectories. The figure depicts the individual differences between real and sham tACS across the four ROIs. Among these, only the superior temporal gyrus shows a significant difference between real and sham tACS

**Fig 5.**
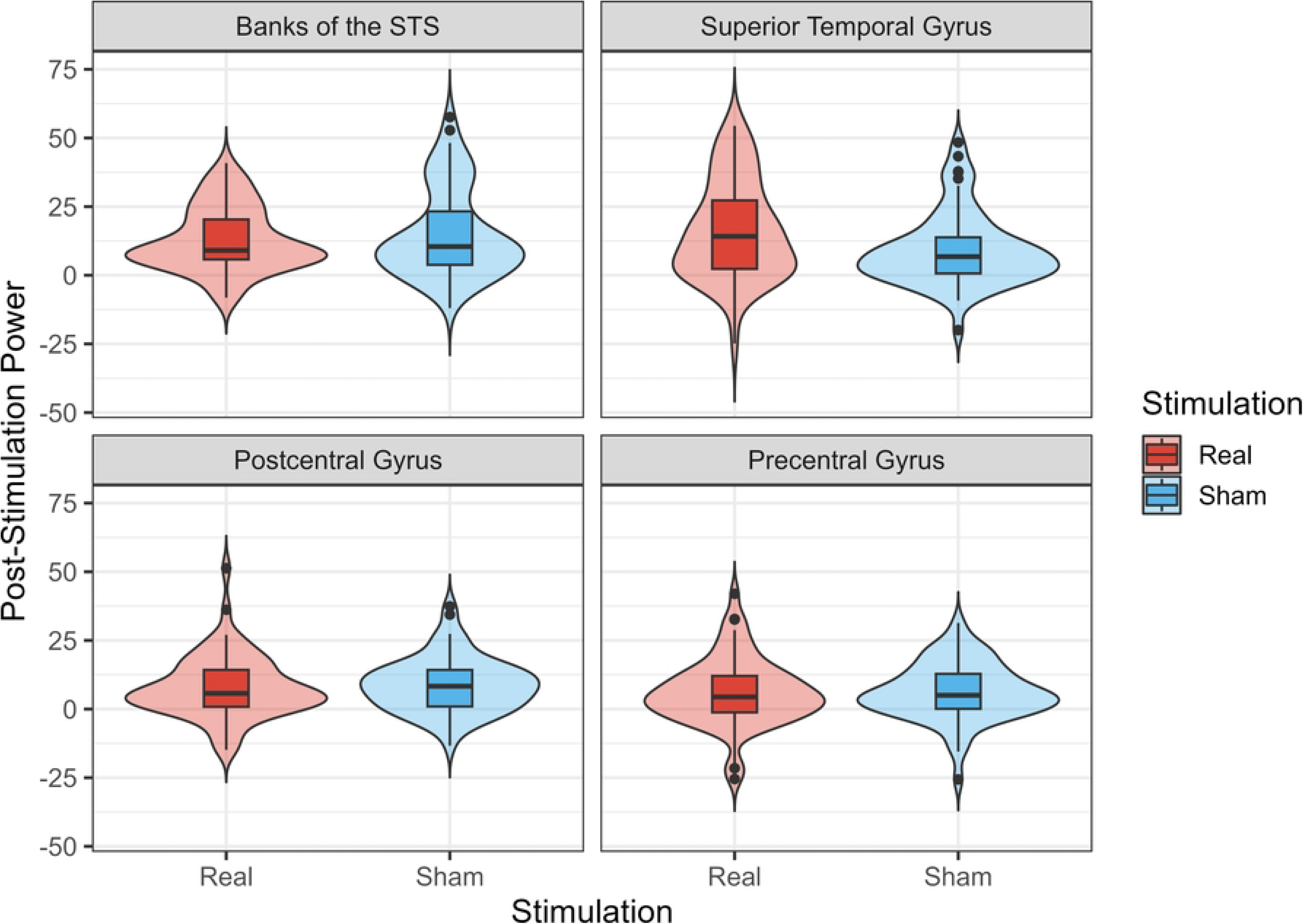
Power distribution by ROI. This figure shows the distribution of gamma power values across conditions (real vs. sham) and the four ROIs. Enhanced post-stimulation gamma power can be selectively found in the STG

To assess whether the effect observed in the STG was driven by one hemisphere, an additional model was conducted. The results revealed a significant main effect of Stimulation (*F*(1, 89.26) = 6.14, *p* = .015), confirming that real tACS was associated with higher gamma power in STG compared to sham stimulation. In contrast, no significant main effect of Laterality was observed (*F*(1,87.47) = 0.11, *p* = .74), nor was there a significant interaction between Laterality and Stimulation (*F*(1,86.65) = 0.21, *p* = .65), suggesting that the effect of stimulation did not differ between the left and right STG.

Further details on the models are provided in the Supplementary Materials.

## 4. Discussion

This study tested whether gamma-tACS could entrain cortical gamma oscillations and whether subsequent 40-Hz auditory stimulation would further enhance the ASSR. According to a state-dependent framework ^42^, pre-modulating cortical activity with tACS should make the cortex more responsive to sensory-driven gamma input.

The primary finding of this study is that real 40-Hz tACS, combined with 40-Hz auditory stimulation, selectively increased the post-stimulation ASSR in the STG compared to sham stimulation. This result was independent of hemispheric laterality and persisted after controlling for pre-stimulation gamma power, indicating that it was not driven by baseline differences in oscillatory activity. The observed enhancement of gamma power in this region is therefore consistent with our hypothesis that combined exogenous (i.e., auditory and electrical) gamma-frequency stimulations can interact synergistically to reinforce oscillatory synchronization within frequency-relevant neural populations. This interpretation is supported theoretical models of oscillatory entrainment proposing that rhythmic external inputs interact with ongoing endogenous oscillations in a frequency-specific manner, preferentially enhancing synchronization in networks that are already tuned to the stimulated frequency ^72^.

Interestingly, tACS electrodes were positioned over bilateral sensorimotor cortices rather than directly over temporal auditory areas. This suggests that the observed modulation of gamma activity in the STG did not arise from direct electrical targeting of the auditory cortex, but rather from network-mediated effects. Anatomical and functional studies have consistently demonstrated strong bidirectional connectivity between SM regions and auditory cortices, particularly auditory-motor integration ^54,55^. Within this distributed network, oscillatory activity in one node can influence distant, functionally coupled nodes ^72–74^. Consistently, stimulation of SM cortices can modulate gamma activity in auditory regions even without direct temporal cortex stimulation ^53^. This further demonstrates that network-level interactions can enhance gamma activity when tACS frequency matches ongoing sensory-driven oscillation ^73^.

In contrast to the findings in the STG, combining 40-Hz tACS with 40-Hz auditory stimulation did not significantly increase the ASSR in the STS, precentral gyrus, or postcentral gyrus. The absence of the ASSR enhancement in the STS may reflect functional differences between the STS and STG. Although the STS plays an important role in higher-order auditory integration ^62^, it is not considered a primary generator of the ASSR, which is typically strongest in the STG ^52^. While the ASSR is not strictly confined to the STG and can be distributed across multiple cortical regions with several neural generators beyond the auditory cortex ^37^, the response is generally maximal within the auditory cortex, particularly in the STG. In this specific context, the lack of gamma enhancement in the STS, together with direct comparisons across regions. may suggest that synergistic interactions between gamma-tACS and auditory stimulation preferentially occur in cortical regions that exhibit a strong gamma synchronization during the ASSR, such as the STG.

It is worth noting that the absence of a synergistic enhancement of the ASSR in SM areas (i.e., the precentral and postcentral gyri) further suggests that the effects of 40-Hz tACS may depend on the presence of task-relevant oscillatory activity. In the present paradigm, such activity is maximal within the primary auditory cortex during the 40-Hz auditory stimulation, which may explain why modulation was observed in the STG but not in SM regions. Therefore, these null results might support the regional specificity of the observed gamma modulation rather than reflecting a lack of sensitivity in the analysis. If the effects of tACS were primarily driven by nonspecific factors, such as broader neural excitability shifts or widespread amplification of gamma power, one would expect comparable changes across multiple cortical regions, particularly those closest to the electrodes. Instead, the selective enhancement confined to the STG, but not in the STS or sensorimotor areas, suggests that the effects of 40-Hz tACS may depend on the functional properties of the targeted network. This pattern is consistent with the idea that gamma-tACS modulates ongoing network-specific oscillatory activity rather than producing a generalized increase in gamma power across the cortex. Furthermore, this interpretation is further supported by the absence of comparable effects in regions not strongly engaged by the auditory stimulation.

The present study represents one of the first attempts to directly combine gamma-frequency tACS with 40-Hz auditory entrainment to probe potential synergistic effects on ASSR-driven gamma activity. Previous studies that investigated the possibility of modulating the ASSR through the application of tACS mainly focused on adopting experimental designs that targeted alpha oscillations, rather than gamma. As a matter of fact, alpha-range tACS has been shown to both decrease ^75^ and to enhance the ASSR in patients with schizophrenia ^76^. Recent studies applying 40-Hz tACS to modulate the ASSR have reported heterogeneous results, often focusing on online effects or by directly placing tACS electrode over T7 and T8, in correspondence to auditory regions ^77–80^. In addition, recent evidence indicates that tACS efficacy depends on electrode montage and the resulting field strength ^56^. Because the auditory cortex is relatively deep, a direct stimulation may be less effective there, whereas more superficial and accessible regions like the SM cortices allow stronger and more consistent modulation. This supports targeting SM cortices as effective upstream sites for modulating auditory gamma activity with tACS. Further studies are needed to unequivocally determine a standard protocol that leads to an enhancement in the ASSR by applying both 40-Hz tACS and auditory stimulation.

The present work has some limitations that should be considered. First, source estimation was conducted using a standard head model rather than individualized realistic head models derived from participants’ MRI data, which were not available. This choice may have reduced the spatial precision of the source localization. Future studies employing individualized anatomical models could improve spatial accuracy and provide more refined insights into region-specific mechanisms underlying tACS effects.

A second limitation concerns the fact that the study did not investigate online tACS effects, focusing only on post-stimulation mechanisms. This was due to the substantial EEG artifacts induced by tACS stimulation, which currently constrain the reliable assessment of immediate neural responses ^81^. Future research employing advanced artifact correction methods may help overcome this limitation.

Third, the absence of additional control stimulation conditions at different frequencies limits the ability to directly disentangle frequency-specific effects from those related to stimulation in general. More precisely, this tACS protocol was defined using a single stimulation frequency at 40 Hz and a peak-to-peak intensity of 2 mA, with a total duration of 20 minutes. In the future, varying stimulation parameters and including control frequencies will be necessary.

Our results demonstrate a region-specific synergistic interaction between 40-Hz tACS and auditory stimulation at 40 Hz, selectively enhancing the ASSR in the STG. This finding supports frequency-specific, network-mediated models of tACS and suggests that combining neuromodulatory approaches may be more effective than single-modality interventions in modulating gamma oscillations. Importantly, this synergy points to potential translational implications for rehabilitative interventions targeting clinical populations with impaired ASSR, such as individuals with schizophrenia ^35,38,39^, AD ^5,33^, and MCI ^40^.

Disruptions in the ASSR and gamma-band synchronization are linked to cognitive deficits, including impaired memory and attentional processing. Previous studies have shown that gamma-frequency stimulation can restore oscillatory activity, enhance memory performance, and even modulate molecular markers associated with AD pathology in animal models ^15,16^. The present findings extend these observations by demonstrating that combining auditory electrical stimulation at 40 Hz can effectively enhance the ASSR in healthy humans, supporting the feasibility of clinical interventions aimed at restoring impaired gamma activity.

Our results may represent a promising avenue for future research. Longer-term applications of gamma-tACS paired with auditory stimulations at 40 Hz could improve cognitive functions in clinical populations with disrupted gamma oscillations. Future studies should investigate optimal stimulation parameters and assess whether enhancements in the ASSR translate into measurable improvements in both cognitive and behavioral outcomes.

## CRediT authorship contribution statement

**Fabio Masina**: Conceptualization, Software, Formal analysis, Writing - original draft. **Rachele Pezzetta**: Writing - original draft, Software, Formal Analysis. **Nicole Genero**: Writing - original draft, Visualization. **Alessandro Tonin**: Conceptualization, Software. **Giorgio Arcara**: Writing - original draft, Supervision. **Daniela Mapelli**: Writing - original draft, Supervision.

## Ethics Approval

The research presented in this paper has been approved by the local ethics committee. All procedures are in accordance with the Declaration of Helsinki.

## Declaration of competing interest

The authors declare that they have no known competing financial interests or personal relationships that could have appeared to influence the work reported in this paper.

## Acknowledgements

This work is funded by the European Union – NextGenerationEU and by the University of Padua under the 2023 STARS Grants@Unipd programme (PHAROS, PHAse in Real-time OScillations: Enlightening the relationship between brain oscillations and brain states). Special thanks to Anna-Lisa Schuler for programming the 40-Hz auditory stimulation (auditory steady-state response).

## Funding

This research did not receive any specific grant from funding agencies in the public, commercial, or not-for-profit sectors.

## Data Availability Statement

The datasets generated during and/or analyzed during the current study are available in the OSF repository, https://osf.io/gmq34/overview?view_only=a190d4432e7842fca4050b20ee8b499d

## Supporting information

S1 Analysis script.

R scripts used to perform all statistical analyses.

S2 Appendix. PECANS questionnaire.

Full version of the PECANS questionnaire,

## Notes

### Competing Interest Statement

The authors have declared no competing interest.

